# Alpha rhythm subharmonics underlie responsiveness to theta burst stimulation via calcium metaplasticity

**DOI:** 10.1101/2025.06.17.660153

**Authors:** Kevin Kadak, Davide Momi, Zheng Wang, Sorenza P. Bastiaens, Mohammad P. Oveisi, Taha Morshedzadeh, Minarose Ismail, Jan Fousek, John D. Griffiths

## Abstract

Repetitive transcranial magnetic stimulation (rTMS) is a non-invasive technique to modulate brain activity, often used in treating Major Depressive Disorder (MDD) by targeting fronto-limbic circuitry. Despite its clinical utility, optimizing rTMS protocols remains challenging due to the complex and variable effects of stimulation parameter changes on synaptic plasticity. Oscillatory brain activity, measurable via Electroencephalography (EEG), serves as a biomarker for functional circuits and treatment response. To better understand the impact of rTMS on brain oscillations and connectivity, we used computational modeling of corticothalamic circuits to explore the mechanisms of stimulus-induced plasticity. We integrated calcium-dependent plasticity (CaDP) with Bienenstock-Cooper-Munro (BCM) metaplasticity formulations in a neural population model of resting-state EEG. By varying protocol parameters, we simulated iTBS effects on spectral power, synaptic efficacy, and calcium concentrations. Our findings highlight a resonance between theta stimulation and individual resting-state alpha rhythms, enhancing incoming excitatory long-term depression (LTD) and inhibitory long-term potentiation (LTP), leading to corticothalamic feed-forward inhibition (FFI). Induced effects were encapsulated by a weakening of corticothalamic loops and enhancement of intrathalamic loops. This work offers a novel paradigm for individualizing iTBS treatments, provides insights into the neurophysiological basis of clinical responsiveness, and offers a framework with which to derive tailored protocols.

## Introduction

Repetitive Transcranial magnetic stimulation (rTMS) is a non-invasive neuromodulatory tool for inducing functional changes in brain circuitry. As such, it is an effective treatment intervention for several neuropsychiatric disorders [1–4], most notably major depressive disorder (MDD) [5–7]. rTMS-MDD treatments have been substantially refined by the superior efficiency and smaller dosages of intermittent theta-burst stimulation (iTBS) against conventional high-frequency protocols (HF-rTMS), which has highlighted the profound influence of pulse structure, pattern, and volume on induced plasticity effects via long-term potentiation (LTP) and depression (LTD) mechanisms [8–13]. However, basic and clinical iTBS studies continue to report a high degree of heterogeneity in outcomes. Here we present and biologically justify an individualized iTBS paradigm for enhancing plasticity effects using a computational model of resting-state activity and rTMS-induced plasticity.

Resting-state electroencephalography (EEG) measures provide valuable insights into the intrinsic function of large-scale brain circuitry, with alpha oscillations (8-13 Hz) being its dominant spectral signature, believed to arise from time-delayed interactions between cortical and thalamic neural populations [14–17]. The frequency of a person’s unique alpha rhythm, or individual alpha frequency (IAF), is considered a stable neurophysiological trait and has been used for evaluating MDD status, responsiveness, and guiding treatments [15, 18–23]. Recent research has demonstrated the utility of IAF as a means of personalizing rTMS treatments to enhance plasticity effects and clinical outcomes [24]. It is postulated that aligning stimulation frequencies with the dominant rhythm of the targeted corticothalamic circuit (i.e. alpha) could engage resonance with its endogenous activity. This may induce stronger plasticity effects to regulate oscillatory dynamics and improve MDD symptoms with greater efficacy than IAF-agnostic protocols [25–29, 18, 30, 19, 23, 21]. As IAFs are variable between subjects [31], their alignment discrepancy with the canonical 10 Hz HF-rTMS frequency may capture a considerable portion of clinical and experimental response variance, and offer a physiological signature upon which to tailor treatments.

The 5 Hz patterned pulse structure of iTBS is an exact subharmonic of the 10 Hz high-frequency rTMS (HF-rTMS), offering a congruence between the two paradigms. We propose a novel premise that iTBS protocols may be designed to act upon the same underlying synchrony mechanism to predict and enhance plasticity effects, but instead via alignment with the IAF’s first subharmonic 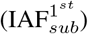. To our knowledge, no research has been conducted to examine iTBS-alpha subharmonic interactions. However, IAF-synchronized HF-rTMS studies have yielded generally promising findings in recent years. A flagship IAF-synchronized rTMS study demonstrated significant sham-controlled improvements in MDD symptoms, with a greater severity of the symptoms predicting stronger outcomes [26, 29, 32]. Further studies have shown and replicated findings that the degree of IAF-rTMS frequency alignment predicted significantly stronger MDD treatment outcomes [18, 19, 23]. Additionally, closed-loop stimulation studies have demonstrated increased neuromodulatory efficacy in phase-locked rTMS interventions targeting frontal IAF oscillations [30] and sensorimotor *µ*-rhythms [33–36]. In TRD patients, IAF phased-locked interventions significantly improved symptoms, but with non-superior efficacy against unsynchronized treatments [37, 38]. Similarly, an earlier study applying rTMS at IAF +1 Hz led to significantly reduced MDD symptoms, although with non-superior efficacy as 10 Hz treatments [39].

Assessing the effects of resonance in any rTMS paradigm remains difficult to probe experimentally, particularly in subcortical brain regions. Computational modelling can address several limitations of empirical EEG measures, as well as challenges posed by the diverse array of rTMS effects. EEG activity may be described and studied mathematically using tractable low-dimensional models of neural population dynamics for isolated examinations of rTMS-induced effects *in silico* [40]. Neural population models are a powerful class of mathematical equations that approximate the activity and properties of neural ensembles with a conservative set of state variables. These models have successfully captured many characteristic brain states and phenomena, including – most relevant to our discussion – eyes-open resting-state EEG regimes [41, 42]. To investigate how rTMS alters neural circuitry underlying resting-state EEG activity, we conducted numerical simulations using a corticothalamic neural population model with calcium-dependent plasticity (CaDP). In this model, rTMS-induced fluctuations in intracellular calcium concentrations determine the direction, magnitude, and timing of plasticity effects[43–46]. These calcium volume changes are homeostatically mediated by a Bienenstock-Cooper-Munro (BCM) metaplastic sliding threshold scheme via n-methyl-d-aspartate receptor (NMDAR) conductances, which ensures that synaptic weights remain both modifiable and stable (Fig. 1) [47–54]. The interplay of these mechanisms accounts for a variety of empirically observed TMS plasticity phenomena, including bidirectionality over pulse dose and time (i.e. LTP to LTD, vice versa)[43, 55–58] depending on paradigm (iTBS vs. cTBS, 10Hz vs. 1Hz). Disentangling these processes may shed light on the high variability in rTMS clinical response rates [59, 60].

**Figure 1:**
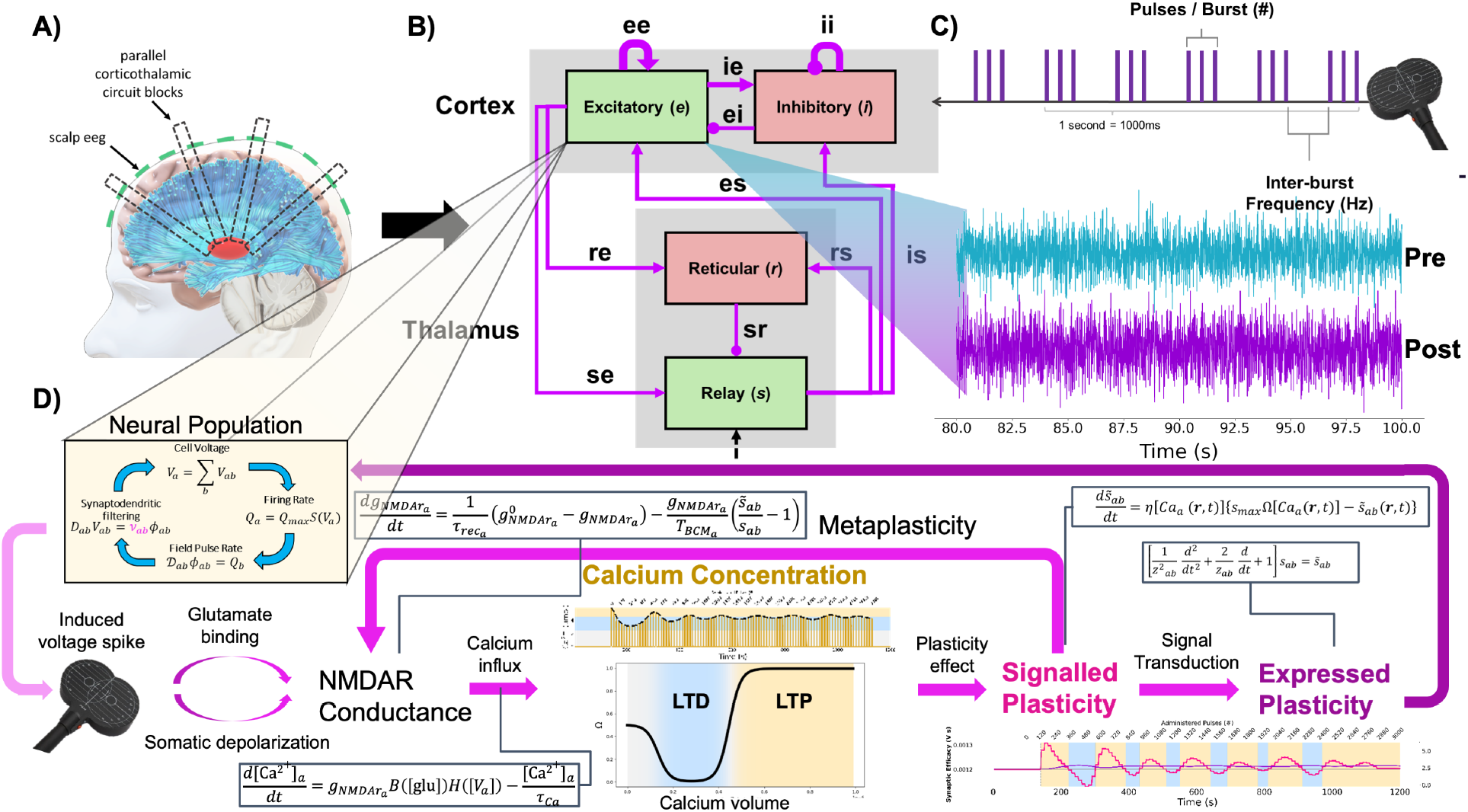
Examining iTBS-induced effects within a computational model of resting-state EEG activity integrated with calcium-dependent plasticity. Schematic of the methodology and neurophysiological model processes. **A)** Corticothalamic column underlying resting-state EEG activity and a **B)** 4-population corticothalamic neural population model of its circuitry, comprised of cortical excitatory pyramidal (*e*) and inhibitory interneuron (*i*) populations, and thalamic excitatory relay (*s*) and inhibitory reticular (*r*) nuclei populations. **C)** Intermittent theta-burst stimulation (iTBS) protocols deliver voltage-spiking stimuli to excitatory cortical populations. Cyan and purple timeseries depict pre- and post-stimulation resting-state activity, respectively, altered by plasticity-induced circuit modifications. **D)** Flowchart of Ca^2+^ scheme wherein post-synaptic depolarization and glutaminergic binding induce metaplastic NMDARs to open, enabling the influx of calcium to drive plasticity effects, with moderate and large concentrations driving LTD and LTP, respectively. Connection weight changes are immediately translated through signalled plasticity (a physiological analog of immediate, though functionally unrealized cell states) before manifesting as fully expressed functional modifications via altered *ν* values, following a signal transduction delay. In conjunction, NMDAR conductance is altered based on the prior weight dynamics, thereby mediating the next instance of calcium influx subsequent plasticity effect. The signaled (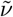; pink) and expressed (*ν*; purple) synaptic weight over pre, active, and post-canonical iTBS are shown for the excitatory-excitatory (*ee*) connection. Yellow and blue windows depict when modulated connection weight is relatively stronger and weaker than initial efficacy, respectively.

Earlier modelling of TBS-induced CaDP with BCM metaplasticity demonstrated that stimuli saturation dynamics and calcium fluctuations can capture the invertible plasticity effects seen empirically [43, 61]. In separate works, alpha resonance was shown to lead to enhanced plasticity effects, and it was speculated that IAF-rTMS may therefore be optimal for treatments [62, 63]. However, these prior works have been confined to cortico-centric representations of the brain and have omitted the role of subcortical structures where rTMS treatment effects are also known to translate [64–66]. The thalamus is an embedded neural structure heavily implicated in alpha rhythmogenesis [17] and a crucially casual hub of MDD pathophysiology that has been increasingly validated as an rTMS treatment target [67, 68, 64, 69, 70]. Abnormal resting-state functional connectivity between the thalamus and frontal cortical structures (i.e. clinical TMS targets) has also been used to distinguish MDD status and predict rTMS treatment success [71, 64]. As rTMS is understood to correct dysfunctional oscillatory synchrony arising from functional imbalances in MDD-implicated pathways (particularly corticothalamic circuitry; [69, 67, 72–74]), spectral modulation may therefore be used as a generalized measure of rTMS effectiveness and a potential brain-based biomarker of clinical responsiveness [75]. Therefore, it is vital to represent the downstream circuitry that engenders EEG activity, is directly implicated in MDD pathophysiology, and plausibly bears deep importance on the interplay between IAF and iTBS.

We investigated how different iTBS protocols affect corticothalamic resting-state power spectra, connection weights, and LTD-LTP calcium levels. Our goal was to understand the relationship between rTMS parameters, individual brain properties (IAF), and resultant plasticity effects to better guide approaches for enhancing induced plasticity in resting-state EEG circuitry. Protocols were varied by pulses-per-burst and inter-burst frequency parameters, which have previously been explored experimentally under the same theorized plasticity mechanisms [55, 76, 77]. Multiple regression analyses (MLR) with a significance threshold of p <.001 were conducted to assess robust effects across distinct alpha frequencies, circuit connections, and loops.

## Results

### iTBS induces resonant modulations to resting-state EEG spectra through alpha rhythm subharmonics

iTBS protocols induced a diverse array of modifications to corticothalamic circuitry, resulting in varying degrees of resting-state broadband power suppression. Power modulation scaled nonlinearly across the stimulation parameter values. Analyzing spectral power changes across stimulation parameter space revealed localized regions where the strongest modulation occurred (Fig. 2A,C). These regions appear close to the natural frequency of the circuit (IAF; 10.5 Hz) and its first subharmonic (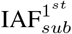 ; 10.5 Hz / 2 = 5.25 Hz). This optimal frequency at the first subharmonic of alpha neighbored the 5 Hz carrier-wave (‘theta’) frequency of canonical iTBS. To confirm that these maximally-modulatory effects occurring in alignment with the alpha rhythm was indeed frequency-dependent, we ran a new set of simulations varying the dominant frequency of the circuit between 8-13 Hz (mimicking variable IAFs in humans) and examined the relative induced modulations. We observed a strong, significant positive correlation between the circuit’s alpha frequency, its first subharmonic, and the inter-burst frequencies of maximally power-suppressing iTBS protocols. Changing the resting-state IAF by modifying the corticothalamic delay parameter *t*_0_ (see section *Corticothalamic resting-state spectra and component measurement*) led to proportional shifts in the stimulation frequencies of maximally-modulating protocols, with the inter-burst frequency of the top modulatory protocols and circuit IAF being strongly correlated (r =.86, p <.001; Fig. 2C). The degree of broadband power suppression induced by canonical iTBS differed over resting-state frequencies, with the strongest modulation occurring at an alpha frequency of 9.5 Hz - wherein canonical iTBS was within 0.5 Hz of its first subharmonic (Fig. 2B, marked green). Consistent with this, we found a significant negative correlation (r = -.54, p <.001) between the simulated iTBS protocols’ distance from the circuit’s resting-state IAF and 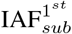-referred hereto as the alpha resonance distance (ARD) - and the degree of broadband power suppression. This negative correlation indicates that protocol-induced power modulation decreased as their inter-burst frequency became increasingly misaligned with the circuit’s alpha frequency (Fig. 2D; top-left). MLR analyses confirmed that this anti-correlation between ARD and broadband power modulation persisted across different alpha frequencies, and was statistically significant across 9 of the 11 unique alpha frequencies following multiple comparison corrections (Table 1). Power modulation effects were also examined across pulses-per-burst, inter-burst frequency, and pulse rate protocol parameters. Significant negative and positive correlations with broadband power were observed, respectively, with the pulses-per-burst (r = -.54, p <.001) and inter-burst frequency (r =.44, p <.001) parameters, but not with pulse rate (r = -.131, p =.048; Fig. 2D). Heightened power modulation reflecting IAF-resonant engagement was visible along inter-burst frequency and pulse rate parameters.

**Figure 2:**
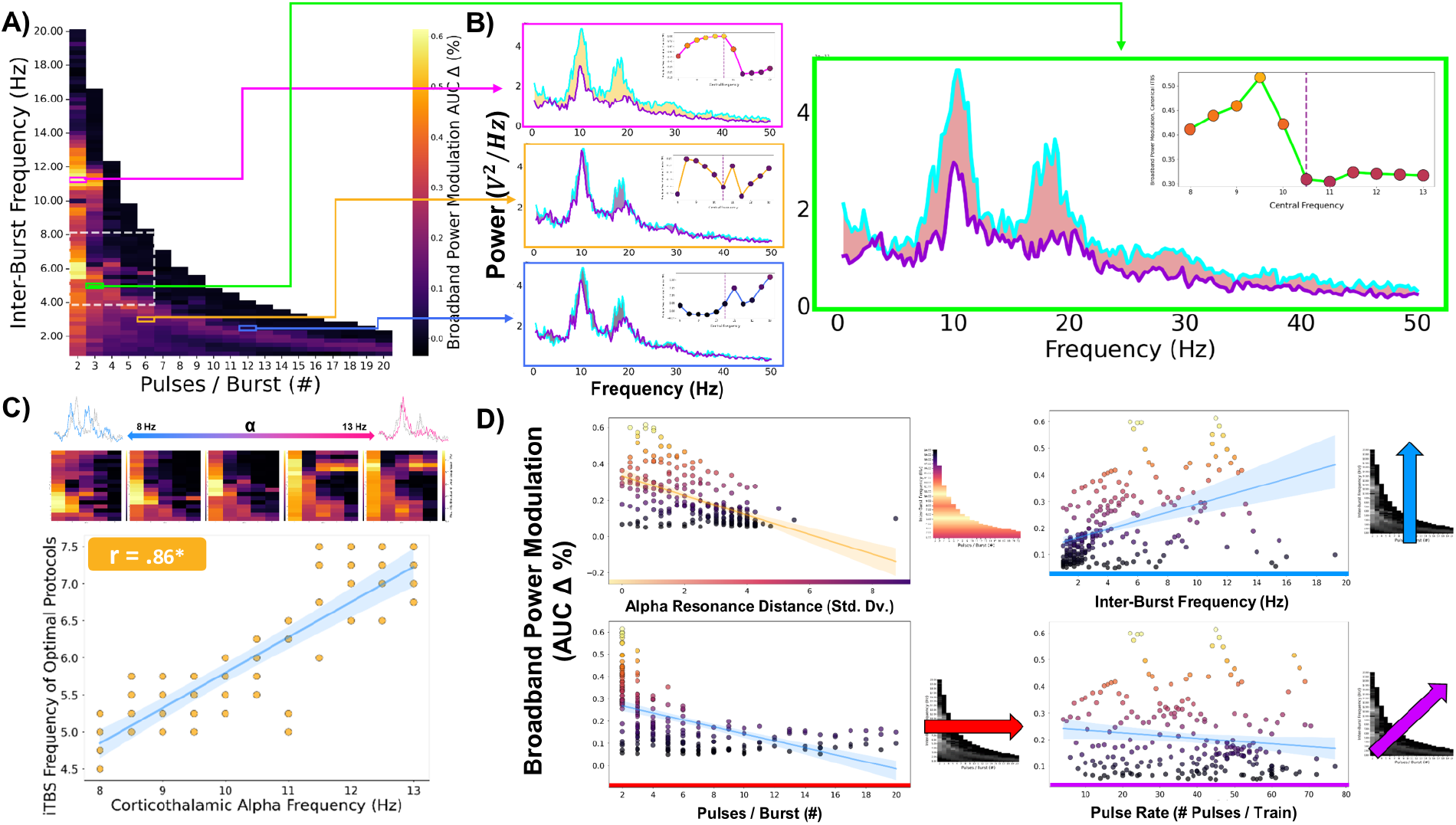
Induced broadband power modulation is enhanced by IAF-iTBS alignment. **A)** Heatmap of iTBS-induced broadband power suppression (pre - post AUC Δ) across parameter space (brighter = stronger suppression), with **B)** samples of power spectra prior to (cyan) and following (purple) stimulation, showing greater degrees of power modulation near alpha harmonic frequencies; green marks the canonical iTBS protocol. **C)** Further simulations with slowing/accelerating the alpha-band frequency show that IAF-iTBS resonance leads to proportional shifts in the inter-burst frequency of the most effective protocols. **D)** Broadband power change as a function of ARD (top-left), pulses-per-burst (bottom-left), inter-burst frequency (top-right), and pulse rate (bottom-right). Together these results support the premise that natural circuit frequency predicts the degree of iTBS-induced broadband power modulation.

### Alpha-resonant iTBS protocols suppress broadband power by enhancing corticothalamic feed-forward inhibition

To better understand the circuit modifications responsible for engendering the broadband power modulations observed in our model, we next analyzed changes in plastic connection weights across the various stimulation parameters. This was done by examining both the directionality and absolute value of pre-vs. post-stimulation change in individual and circuit-mean connection weights (see Supplemental Fig. S1 for individual weight change heatmaps). As with the power modulations shown in Fig. 2, circuit-mean synaptic weight change showed a nonlinear, non-monotonic relationship with stimulation parameters (Fig.3A, C). Interestingly, we found that different iTBS protocols induced synaptic plasticity in unique ways to circuit connections, varying in both the number of affected connections and the magnitude of their modification. A sample of radar plot profiles shown in Fig. 3B demonstrate the unique weight change patterns induced by different protocols. A significant correlation between circuit-mean synaptic weight change was observed for inter-burst frequency (r = -.31, p <.001) and pulse rate (which showed a Gaussian distribution of modifications), but not for pulses-per-burst (Fig. 3C). ARD also showed no relationship with circuit-mean synaptic weight changes, indicating that IAF-iTBS resonance did not influence the degree of the cumulative plasticity effect. Notably, there was no robust relationship between circuit-mean synaptic weight change and broadband power modulation (r =.196, p =.003). There was also considerable heteroscedasticity in the broadband power modulation values, with a progressively larger variance at higher levels of circuit-mean absolute synaptic weight change (3D). This result shows that circuit-wide synaptic weight modification alone is not a robust predictor of broadband power modulation, since large synaptic modifications can result in either low or high extents of power modulation. Next, we looked closer at individual connection weights within the circuit. We refer to connections whose values became larger relative to their pre-stimulation weight as *enhanced* (LTP; larger in magnitude and further from zero) and smaller as *dampened* (LTD; smaller in magnitude and closer to zero). MLR analyses showed significant relationships between synaptic weight change and broadband power modulation for several connections, with weight enhancements occurring within afferent connections to cortical inhibitory (*i*) and thalamic reticular (*r*) populations, and dampening within cortical excitatory (but not thalamic relay, *s*) populations. All inhibitory-afferent connections (i.e. inputs to inhibitory populations) demonstrated a significant interplay with ARD (Fig. 3E). However, only a subset of these connections were significantly modified as a function of ARD. Thalamic reticular-afferent connection weight change was negatively correlated with ARD, while cortical excitatory-afferent weight change was positively correlated, implying that ARD led to enhancement of thalamic reticular-afferent weights and dampening of cortical-afferent weights. These results are outlined in Table 2. This general phenomenon, whereby inhibitory activity is enhanced via excitatory inputs onto inhibitory neurons, is known as feed-forward inhibition (FFI). Thus, our results indicate that alpha-resonant iTBS protocols achieve broadband power suppression via selective modulation of corticothalamic FFI.

**Figure 3:**
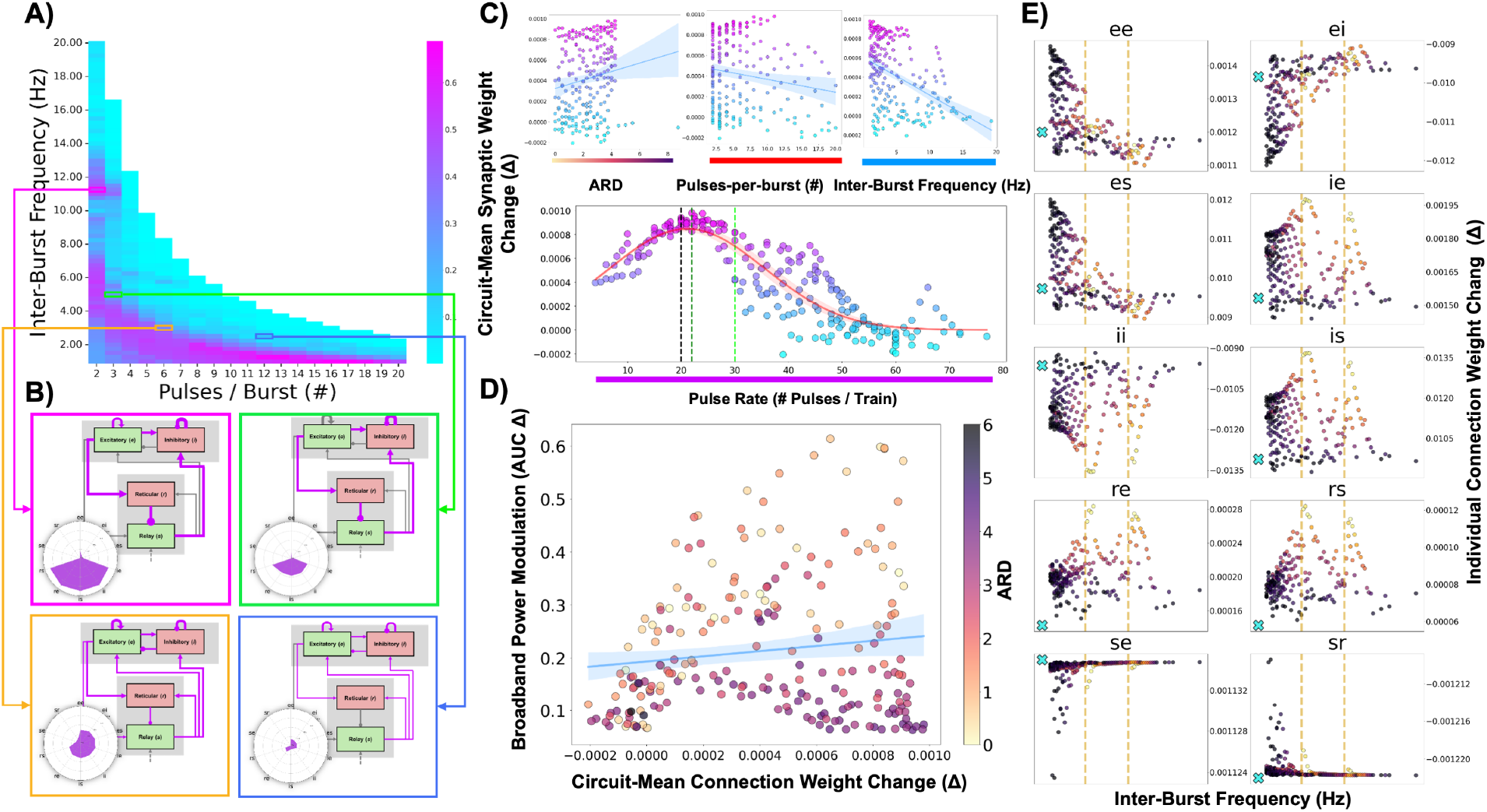
Selective synaptic modifications underlie broadband power modulation and demarcate protocol efficacy from circuit-wide plasticity effect. **A)** Heatmap of circuit-mean connection weight changes revealed a confined region of enhanced modification across parameter space distinct from broadband power suppression. **B)** Connection modification patterns for a sample of protocols illustrate unique synaptic plasticity effects from generalized circuit weight changes. **C)** Circuit-mean modifications adhere to a Gaussian distribution over protocol pulse rate, highlighting an optimal range within which plasticity is induced. Moreover, **D)** group-level weight modification and broadband power modulation are not statistically associated, implying a necessity to induce selective plasticity effects to compel functionally efficacious changes. **E)** Selective modifications are driven further by resonant effects, which visibly enhance inhibitory-afferent LTP across circuit connections.

### Upregulation of feed-forward inhibition is corroborated by resonant engagement selectively facilitating and attenuating LTP-LTD calcium influx

Calcium release was induced during active iTBS across circuit connections, with different calcium volumes evoking different plasticity effects (LTD, LTP). As each plastic modulation altered NMDAR conductance rates, induced calcium volumes evolved non-monotonically over delivered pulses. To probe the low-level physiological drivers underpinning connection modifications across the circuit, and how they were influenced by rTMS pattern, we examined the relationship of LTD-LTP calcium volumes on broadband power and connection weight change, and as a product of ARD and pulse rate, the results of which are outlined in Table 3. Consistent with connection weight changes, LTP calcium volumes in circuit-wide inhibitory-afferent connections and LTD volumes in cortical excitatory-afferent connections were robustly associated with broadband power modulation. This implies that iTBS treatments drive FFI through calcium innervation, and IAF-resonance amplifies these influx patterns. Only circuit-mean LTP volumes, and not LTD volumes were significantly correlated with synaptic weight change (r =.84, p <.001) (Fig. 4E, F), implying that rTMS-induced LTP calcium was the principal driver of system change. Next, frequency interactions on calcium volumes were examined. MLR analyses were performed to compare ARD and induced LTD and LTP calcium volumes in each connection of the circuit. Thalamic relay-afferent connections *se* and *sr* were not included in these comparisons as LTP calcium volumes were never induced within them. Interestingly, protocol ARD predicted systematically increased LTP calcium levels in thalamic inhibitory-afferent connections but attenuated levels in cortical excitatory-afferent connections, implying that resonant engagement facilitates influx for the former and blocks it for the latter. The opposite was true of LTD calcium, whose levels showed inverted trends with ARD, but were narrowly non-significant following multiple comparison corrections. Resonance-driven LTP calcium levels in thalamic *rs* connections and attenuated levels in cortical *ei* and *es* connections were statistically significant. Pulse rate was moderately and strongly anti-correlated with circuit-mean concentrations of LTD (r = -.36, p <.001) and LTP (r = -.85, p<.001), respectively, meaning that generalized calcium levels were attenuated in faster protocols (Fig. 4E, F).

**Figure 4:**
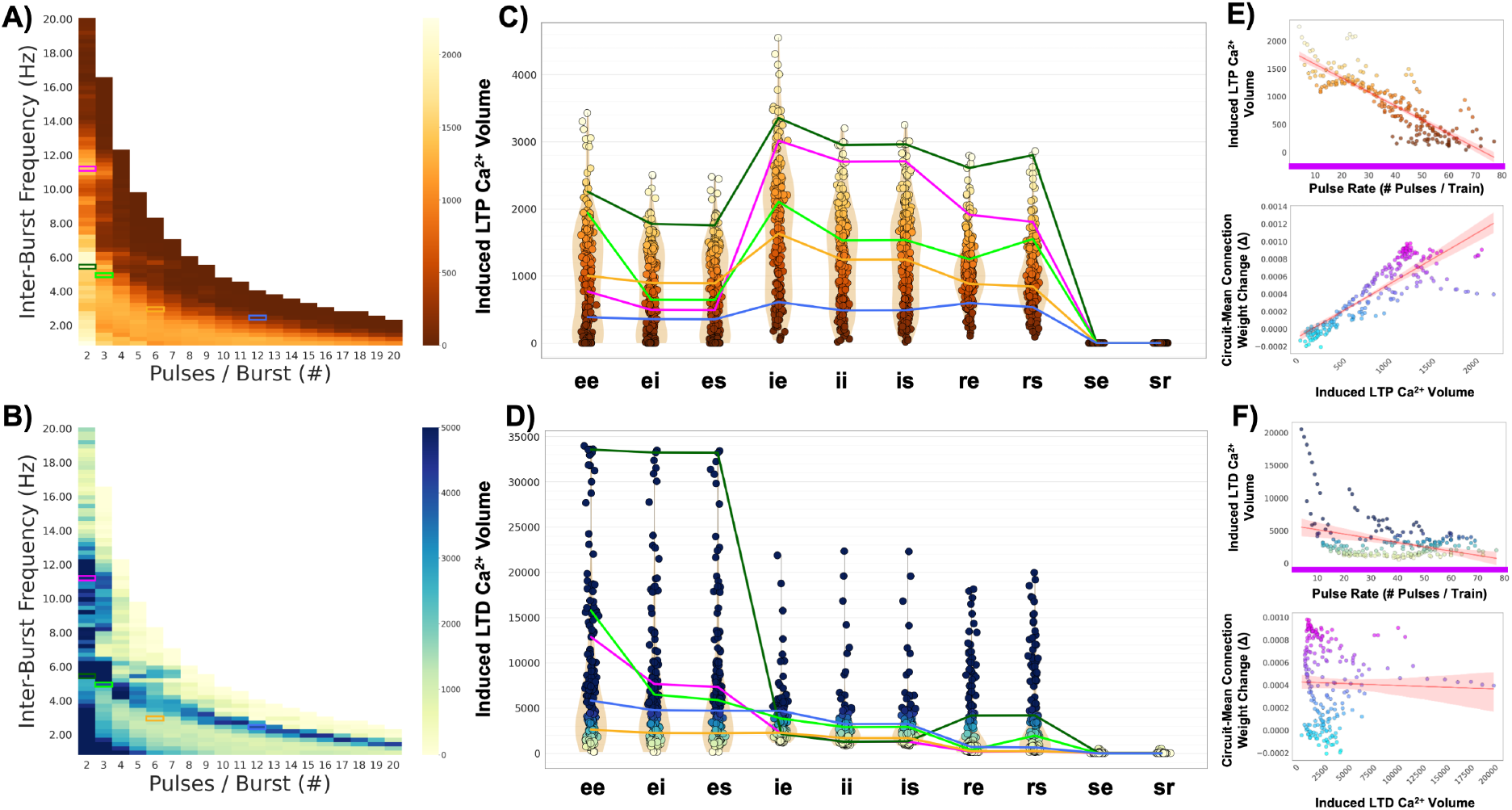
Induced calcium concentrations corroborate a two-pronged effect on spectra via inhibitory LTP and excitatory LTD. Heatmaps of circuit-mean **A)** LTP and **B)** LTD calcium volumes, as well as **C)** LTP and **D)** LTD volumes for each corticothalamic connection. Parameter changes induced unique patterns of calcium release, with protocol efficacy being defined by an amplification of LTP calcium globally and LTD calcium within cortical-excitatory afferent connections. These unique calcium effects are demonstrated across subset of different protocols, including standard (lime green) and model-optimal (dark green) iTBS. **E)** Circuit-mean LTP calcium volume decrease was strongly predicted by pulse rate increase and strongly predictive of circuit-mean connection weight change, while **F)** LTD volume decrease was only moderately predicted by pulse rate increase and not predictive of weight change.

### Stimulation responsiveness is encapsulated by corticothalamic dampening and intrathalamic enhancement

To understand rTMS-induced effects on the circuit macroscopically, connection gains were computed (Eq. 14) and aggregated into the system’s three major feedback loops (schematically depicted in Fig. 5A): Cortical (X), Corticothalamic (Y), and Intrathalamic (Z) (Eqs. 15, 16, 17). This enabled pre- and post-stimulation circuit states to be projected into the 3D XYZ loop gain space, which has been shown to capture all major dynamical regimes of the Robinson corticothalamic model [41, 78–82]. Our analyses showed that different iTBS protocols induced varying displacement in XYZ space from the initial pre-stimulation loop gains. These are shown in Fig. 5B, where each point in the space corresponds to a post-stimulation loop gain configuration induced by a unique protocol. MLR analyses (Table 4) revealed a modest positive influence on broadband power modulation from X loop gains (t = 4.18, p <.001), a strong negative influence from Y loop gain (t = -23.05, p <.001), and a strong positive influence from Z loop gains (t = 25.46, p <.001) (Fig. 5C). Circuit modulation was thus strongly predicted by a protocol’s propensity to dampen corticothalamic loop weights and enhance intrathalamic loop weights. A separate MLR analysis was performed to examine the resonance engagement of each loop gain. Significant resonant interactions occurred in cortiothalamic and intrathalamic loops, with ARD showing positive association with Y loop gains (t = 5.74, p <.001), negative association with Z loop gains (t = -8.22, p <.001), and no effect on X (cortical loop) gains (Fig. 5C, D). Modulatory resonance enhancements were also apparent across iTBS parameters in Y and Z, but not X loop gains (Supplemental Fig. S2).

**Figure 5:**
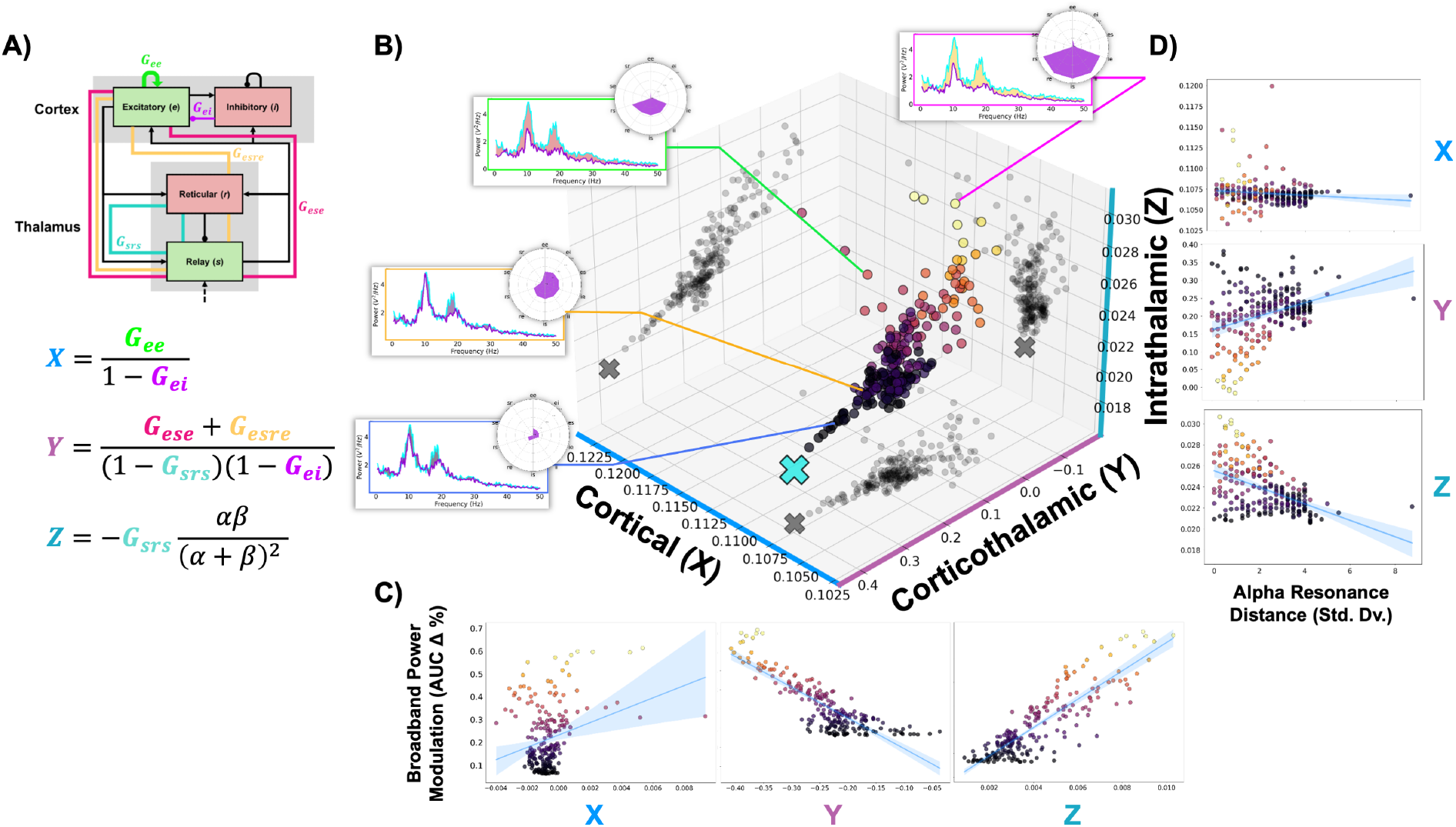
iTBS efficacy is robustly captured by corticothalamic dampening and intrathalamic enhancements of circuit loop gains. **A)** Corticothalamic connection gains may be mathematically collapsed into the circuit’s three major loop gains: X (Cortical), Y (Corticothalamic), and Z (Intrathalamic). **B)** Post-stimulation circuit distributions in XYZ loop gain space show a robust trajectory in relation to broadband power change, with canonical (green) and a subset of novel iTBS protocols induce different displacements in space, corresponding to unique neurophysiological modifications. Point colors reflect broadband power modulation (as seen in Fig. 2A). **C)** Broadband power changes are shown to be a robust product of decreases and increases to Y and Z loop gains, respectively, implying downstream modifications bear the most functional importance on broadband power changes. These trends are further amplified by **D)** alpha resonance distance, which also manifests downstream to more strongly displace Y and Z circuit loop gains.

## Discussion

We modeled and examined rTMS-induced plasticity effects on broadband power, synaptic weights, LTP/LTD calcium, and circuit loop gains within a corticothalamic neural mass model of resting-state EEG activity. We observed pattern-dependent modulation of these components, which were further amplified by a resonant interplay between IAF subharmonics and iTBS frequency. Here we outline the implications of these induced effects and outline the brain-stimulation mechanisms underlying them.

### Mechanisms of action underlying iTBS protocol efficacy

Previous empirical and modelling studies have examined stimulation effects across different amplitudes [83, 84, 59], pulse dosages [85–87], inter-train intervals [88, 89], treatment lengths [90, 91], session intervals [92], and other parameters against physiological and clinical outcomes. Here we extend this research by examining how stimulus pattern affects induced plasticity and their relationship to resting-state spectra. We propose that the dynamics observed in modelled parameter space, alongside empirically documented theta-burst effects, reflect the balanced engagement of four pattern-based mechanisms through which iTBS induces plasticity and drives robust responsiveness:

1. **Endogenous theta mimicry:** Theta oscillations are strongly implicated in active neural encoding processes, including learning and memory, and was the basis upon which the theta-burst stimulation protocol was originally formulated and adapted for human interventions [93, 94]. TBS mimics the frequencies associated with endogenous theta rhythms, which play a major role in plasticity phenomena in hippocampal tissue. This more effectively engages the mechanisms of NMDAR activation, calcium influx, and subsequent signaling cascades for LTD-LTP to achieve comparable plasticity effects as unpatterned rTMS with smaller pulse doses.
2. **Intermittency:** The intermittency of the iTBS pulse train structure allows for a larger calcium influx between recovery periods necessary for nominal LTP effects. This is in contrast to the continuous structure of cTBS which drives a consistent moderate calcium influx associated with LTD, in line with the principles of the calcium control hypothesis [44].
3. **Pulse rate:** Within the intermittent paradigm, the rate with which canonical iTBS delivers pulses lands within a range suitable for driving calcium innervation without oversaturating and downregulating the permeability of NMDARs mediating their plasticity effects, compelling more efficient and robust effects than protocols with smaller pulse intervals.
4. **Circuit resonance:** When delivered at a balanced intensity, the 5 Hz theta frequency of canonical iTBS enhances calcium innervation and subsequent LTP effects with greater efficacy when more closely aligned with the IAF (first subharmonic), which naturally varies between individuals.

Mechanisms 1 and 2 are well-established in literature [95, 96, 50, 93, 97], and our results strongly corroborate these proposed mechanisms of pulse rate (3) and circuit resonance (4) in several ways. Regarding pulse rate, we found an evident disadvantage for faster protocols, which venture away from the established theta-burst regime, have smaller

recovery periods, and increase the circuit’s susceptibility to stimulation oversaturation compared to slower protocols, all of which attenuate LTP calcium - the driving force underlying efficacious circuit-wide modifications. Broadband power modulation was maximized in protocols with lower pulses-per-burst and inter-burst frequencies aligned with the first subharmonic, rather than the fundamental alpha (Fig. 2A). Pulse rate also had no predictive effect on broadband power change, implying no modulatory advantage for faster protocols (Fig. 2D). These effects were also present in fixed-amplitude conditions (Fig. S6). Notably, circuit-mean synaptic weight change was normally distributed across stimuli pulse rate, with the model-optimal protocol (2 pulses-per-burst, 5.50 Hz inter-burst frequency; 22 pulse rate) being congruent with the ideal pulse rate and neighboring that of HF-rTMS (20 pulse rate), as exemplified in Supplemental Fig. S4. Saturation effects were exemplified in fixed-amplitude protocol simulations, which showed that higher pulse rates strongly predicted increases in NMDAR modulation, thereby leading to metaplastic mechanisms being engaged and overall decreases in LTP calcium levels and broadband power modulation (see *Scaled vs. fixed metaplastic effects*). Finally, LTP calcium was strongly anti-correlated with pulse rate (Fig. 4E), suggesting that, even following amplitude-scaling, slower protocols compel the most favourable and robust modifications.

Our results also strongly emphasized the presence and modulatory enhancement of circuit resonance. We found robust relationships between protocol ARD and broadband power modulation (Fig. 2D), enhanced LTP calcium levels within thalamic reticular-afferent connections (Fig. 4A) and loop gains (Fig. 5D), and circuit-wide FFI. Resonant interactions were validated by proportional shifts in the efficacy of iTBS frequency across corticothalamic alpha frequencies (Fig. 2C). Protocol efficacy was characterized by enhanced LTP calcium in inhibitory-afferent projections and simultaneous LTD calcium in cortical excitatory-afferent projections. It was further revealed that ARD predicted an attenuation of opposing calcium concentrations in a subset of these connections, revealing an elegant two-pronged mechanism that drove and reinforced FFI (Fig. 4). This is especially notable given the diverse modification patterns observed across parameter space, and aligns with studies suggesting different iTBS protocols can uniquely affect cortical inhibition [98].

Anatomically speaking, the robust modulation of broadband spectral power observed in our model was driven by changes in corticothalamic and intrathalamic loops that lie downstream of the superficial cortical site where stimulation was applied. This finding is consistent with previous work showing that rTMS-TRD treatment success [67] is better predicted by stimulation-induced changes to fronto-thalamic, rather than frontal brain connections. Enhanced functional connectivity in corticothalamic circuits has also been identified as a core pathophysiological feature of MDD [71, 99], with symptom severity correlating with degree of hyperconnectivity [69]. Resonance manifesting in downstream corticothalamic and intrathalamic loop gains, but not in superficial cortical loop gains, corroborates the mechanism of action proposed by Leuchter and colleagues that IAF-rTMS may reset thalamocortical oscillations more effectively than frequency-agnostic protocols to correct aberrant functional configurations and drive clinical improvements [100, 28, 14, 27, 101, 18]. This resonance mechanism has clinical relevance given that heightened alpha power is a distinguishing EEG feature of MDD [102, 103, 20, 104, 105, 22], and its suppression has been linked to treatment responsiveness [106–109]. Studies have shown that individuals with more severe baseline symptoms of depression and anxiety showed enhanced responsiveness to alpha-synchronized rTMS [32, 29], suggesting that IAF-rTMS may produce selective therapeutic effects for neuropsychiatric disorders characterized by abnormal resting-state patterns. Supplemental modelling analyses further revealed that alpha-band power suppression was strongly correlated with broadband power suppression and X and Y loop gain changes (Supplemental Fig. S3), supporting that rTMS also acts on alpha power via downstream circuit modifications. This is consistent with the dynamical regimes of the XYZ space as the distribution of induced loop gain weight changes captures a trajectory that departs from the alpha regime, leading to a reduction in power [42, 81].

We posit that iTBS leverages these four mechanisms over HF-rTMS to maximize nominal LTP effects and could explain why HF-rTMS treatments take longer to induce a comparable plasticity effect [110, 8]: A uniform, high-frequency pulse pattern and lack of an OFF period over-engage the homeostatic stability mechanisms described by BCM theory, leading to decreased NMDAR conductance despite administering pulses at similar rates. The clinical relevance of this mechanism is supported by empirical results showing that MDD patients have circuits with preferred resonant frequencies, and those with stronger resonance engagement at 10 Hz demonstrated both greater symptom severity at baseline and showed greater improvement with 10 Hz rTMS treatment than other slower or faster alpha frequencies [111]. Placebo-controlled 5 Hz rTMS has shown clinical effectiveness and non-inferior efficacy to 10 Hz treatments [112–119]. However, IAF has only been found to be predictive of improvement for 10 Hz treatments thus far [18]. This may be because 5 Hz rTMS uses only half the pulse rate of 10 Hz, suggesting that more stimuli are necessary to meaningfully engage circuitry before IAF-resonance becomes physiologically relevant. This suggests that resonance may be engaged via the IAF, but that protocols must remain within an optimal pulse rate range to qualify their induced effects, implicating both premises 3 and 4.

### Individualized iTBS treatments, clinical applications, and takeaways

Several methodological considerations warrant discussion. Our model assumes equality in the neuromodulatory capacity of each TMS pulse and LTD/LTP calcium thresholds across circuit connections, which may oversimplify the biological mechanisms underlying rTMS-induced plasticity effects. We address this by scaling TMS pulse amplitudes in our primary analyses emulate neural adaptation effects and isolate the contributions of stimulation pattern and saturation from protocol intensity. In addition, we assumed uniform LTD and LTP calcium thresholds across all circuit connections, which may be more accurately represented on an individual basis for calcium dynamics within each neural population or across circuits.

Whilst our primary findings (i.e. IAF resonance and low-intensity efficacy) and most of our results were consistent between scaled- and fixed-amplitude stimulation, some disparities still emerged, and these differences have implications for factors such as the choice of intensity ranges in physiological experiments. Thus, our approach enabled large-sweeping explorations of the rTMS parameter space that would not be possible without stimulation amplitude scaling, and future research should examine these assumptions in detail.

Key areas for future investigation include understanding the modulatory drop-off between fundamental and subharmonic alpha resonance, exploring individual power spectra components’ interactions with iTBS, investigating spatial dynamics and stimulation targets [20, 120], and examining other temporal components such as intra-burst frequency [121]. Fitting individualized plasticity effect timescales, rather than using fixed values, could help explain response variability [43].

IAF-guided iTBS presents a promising approach for individualizing MDD treatments while requiring only minimal protocol modifications. Our results showed that resonant protocols targeting the first subharmonic of resting-state alpha induced the strongest modulation of resting-state spectra through excitatory-afferent LTD and inhibitory-afferent LTP calcium dynamics. These changes manifested in the dampening of corticothalamic loop gains and enhancement of intrathalamic gains.

Because alpha peak centrality is a robust and stable trait, patients’ IAFs can be determined from a single resting-state EEG measure, even using low-cost mobile systems [15, 122]. We provide a neurophysiological basis for rTMS response and a computational framework for deriving individualized protocols. We hypothesize plasticity effects to undergo significantly greater modulation via IAF subharmonic resonance and encourage researchers to examine frequency alignment in iTBS datasets as a predictor variable, as our findings have important implications for understanding MDD mechanisms and improving treatment efficacy through personalized protocols.

## Methods

The present work employed 4-population corticothalamic neural model, composed of excitatory pyramidal *e* and inhibitory interneuron *i* populations of the cortex, as well as excitatory relay *s* and inhibitory reticular *r* nuclei populations of the thalamus. Analyses examined relative changes in baseline synaptic weights and excitatory resting-state spectral power following 600 pulses of iTBS administered to the excitatory cortical population *e*. Primary analyses had all iTBS protocols administer pulses at an amplitude proportional to their stimulation intensity, and at a fixed amplitude (see *Stimulation intensity and protocol amplitude scaling*), delivered at a 50 Hz intra-burst frequency (20 ms pulse intervals) for 2 seconds ON, followed by 8 seconds OFF [95]. Stimulation protocols were varied across iTBS ‘pulses-per-burst burst (#)’ and ‘inter-burst frequency (Hz)’ parameter dimensions.

rTMS and iTBS have been shown to induce a diverse set of bidirectional spectral modulations across an array of frequency bins, and different protocol patterns may translate within unique components of the spectra [104, 123, 124]. Thus, to capture this range of effects, we examined broadband power modulation as the primary outcome and coefficient of rTMS-induced modification.

### Neural population models

Neural population models are a set of mathematical equations that approximate the firing dynamics and physiological properties of neuronal populations to describe meso-macro scale activity in the brain. We refer the reader to [42] for a holistic explanation. In a cyclical process, the mean somatic potential *V*_*a*_ of type *a* neurons are the result of summed afferent inputs from type *b* populations, captured as

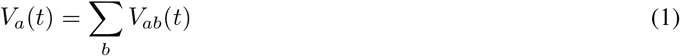

where *V*_*ab*_ represents the post-synaptic potential of the recipient population *a* as a product of incoming inputs from population *b* at time *t*. Type *b* populations may represent an ‘endogenous’ population within the circuit (ie. *e, i, r*, or *s*) or an external stimulus input (ie. *x*; TMS). The synaptodendritic dynamics which mediate incoming inputs are described as

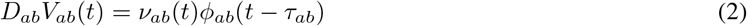

where coupling strength *ν*_*ab*_(*t*) is the product of the mean synaptic weight *s*_*ab*_ multiplied by the mean number of synaptic connections *N*_*ab*_ between *a* and *b* populations (which we do not modify here). *ϕ*_*ab*_ is the mean axonal pulse rate from *b* to *a* and *τ*_*ab*_ is its signal propagation time delay between the populations; due to, for instance, the spatially extended pathways between cortical and thalamic populations. The operator *D*_*ab*_ describes the time evolution of *V*_*ab*_ in response to synaptic input, given by

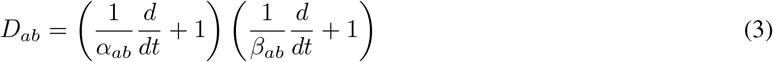

where *α*_*ab*_ and *β*_*ab*_ describe the inverse decay and rise response rates of the post-synaptic potentials, respectively. The resulting mean firing rate *Q*_*a*_ is related to mean somatic potential *V*_*a*_ through a sigmoidal function *S*_*a*_, defined as

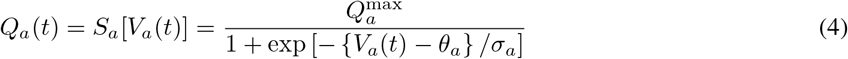

where 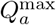 is the maximum possible firing rate, *θ*_*a*_ is the mean firing threshold, and *σ*_*a*_ is the standard deviation of the threshold. Finally, the firing of population *a* neurons generates axonal pulses which propagate along axons toward novel synapses of population *c* neurons. The mean axonal pulse rate is related to the mean firing rate by

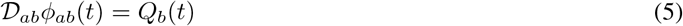

where

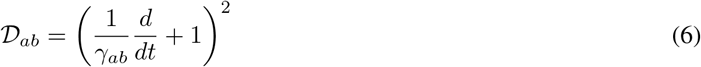

The parameter *γ*_*ab*_ represents the axonal propagation rate from neurons of type *b* to type *a*.

### Calcium-dependent metaplasticity

We outline the approach of [40, 43] and reiterated by [61, 125] employed in the current work. A corticothalamic population model was outfitted with CaDP formulations incorporating a metaplastic Bienenstock-Cooper-Munro (BCM) sliding threshold scheme. Here, metaplasticity describes the dynamics of synaptic connections within the model whose weights ultimately change as a product of post-synaptic intracellular calcium (Ca^2+^) volumes, which themselves are modulated by higher-order NMDAR calcium permeability rates (hence ‘meta’) [44, 45]. Synaptic weight values between an afferent *a* and efferent *b* population are plastic and dynamically modulated via the synaptodendtritic coupling strength term *ν*_*ab*_, altered through its mean synaptic weight term *s*_*ab*_, with expressed synaptic weight formulated as

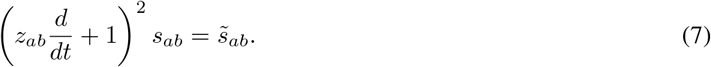

Physiologically realized changes to synaptic weights are hypothesized to transpire following a protein cascade transduction from sporadic calcium-medicated plasticity processes. The given instance when connection weights are modified that eventually translate to measurable changes in synaptic efficacy, known as signaled synaptic strength 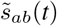 and modelled as

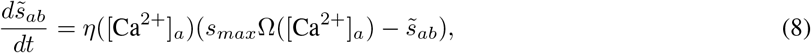

allows for non-monotonic changes in synaptic efficacy over time, leading to either functional LTP or LTD depending on the instance when stimulation is halted and modification concluded. As such, expressed synaptic weight acts as a low-pass filtered, homeostatically-stable translation of signalled synaptic weight change that will converge towards its modified end-value, with *z*_*ab*_ being the response timescale of the transduction delay between signalled and expressed plasticity. Put simply, signalled plasticity represents the immediate physiological synaptic efficacy state that is not yet functionally manifest while expressed plasticity represents the time-delayed, realized weight of the synapse that converges towards modified signalling. Post-synaptic calcium ion concentrations Ca^2+^ determine the directionality (LTP/D) and magnitude of induced plasticity at the point when stimulation is halted. Synaptic weight modification is driven by the calcium control functions of the plasticity rate

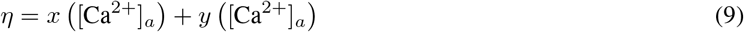

and the determinant plasticity effect induced

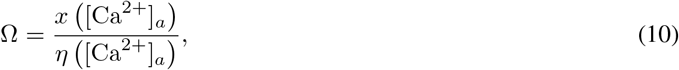

where low calcium concentrations lead to no meaningful plasticity effect, moderate concentrations induce LTD, and high concentrations induce LTP (to capture the BCM activation function). *x* and *y* are the calcium-dependent potentiation and depression rate functions, respectively, ascertained empirically from Shouval et al. [44, 45] (see [40] for the full expansion). Ca^2+^ is the product of pre-synaptic glutaminergic release and post-synaptic somatic depolarization, both of which are required for NMDARs to open and allow ionic influx, captured as

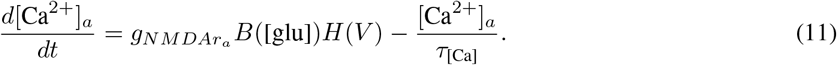

*g*_*NMDAr*_ is the NMDAR conductance-mediated calcium permeability, *B* is the extent of NMDAR glutaminergic binding based on a sigmoid function of glutamate concentration [glu], *H* is the voltage-dependent threshold for NMDAR modulation (a nonlinear function which, increases with *V*, except for exceedingly strong depolarizations), and *τ*_[Ca]_ is the timescale of calcium dynamics - the frequency with which calcium volumes update. Metaplasticity dynamics are integrated as a BCM sliding threshold scheme to describe adaptive changes in the *g*_*NMDAr*_ term, which would be otherwise fixed with base CaDP formulations (as seen in [40]). Here, the NMDAR conductance *g*_*NMDAr*_ that regulates post-synaptic Ca^2+^ influx rates and delineates LTD-LTP induction is moderated on the prior states of synaptic efficacy *s*_*ab*_.

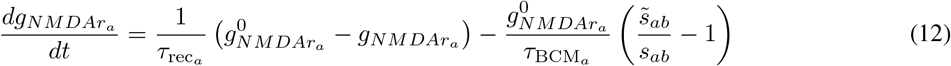

where 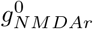 is the calcium conductance at stability, *s*_*ab*_ is the expressed synaptic weight, and 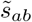 is the projected synaptic weight towards which it trends. *τ*_BCM_ and *τ*_rec_ are the timescales for metaplasticity dynamics and calcium conductance to recover to the stable value, respectively. Post-synaptic glutamate concentrations [glu] are a product of pre-synaptic activity, which decay over a timescale *τ*_[glu]_, modeled as

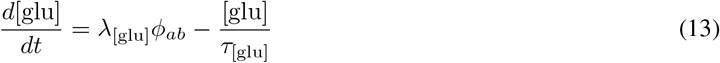

where *λ*_[glu]_ is the mean glutamate concentration released per incoming pre-synaptic excitatory spike, *ϕ*_*ab*_ is the incoming pre-synaptic flux from excitatory populations, and *τ*_[glu]_ is the timescale of glutamate decay.

### Corticothalamic loop gains and space

Prior gain calculations by Robinson et al. have used the steady-state 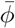 to compute gains. However, because system states are continuously undergoing plasticity-induced changes throughout simulations, here we calculate the gain values at each time step. Altered system states are then ascertained by computing mean gain values from the final 10 seconds of simulated activity.

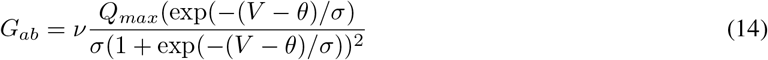

The individual gains for each connection can be reduced into 3 aggregate loop gains that capture the three major loops of the corticothalamic circuit: Cortical (X), Corticothalamic (Y), and Intrathalamic (Z), computed as

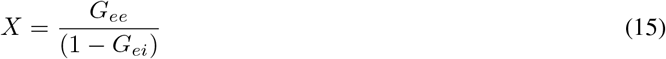

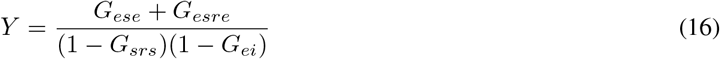

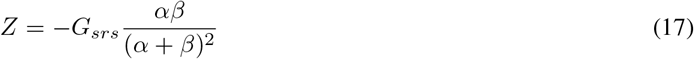

### Model parameters and settings

All model-simulated outputs were generated with a step size (*dt*) of 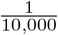 and a write interval of 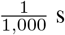.

Corticothalamic state parameters were taken from from the NFTsim.conf library [126], outlined in Table 5. Initial *ee* and *ei* weights were modified within 20% of default value range to maintain circuit stability during and following stimulation, and to retain the oscillating evolution of plasticity signalling in excitatory-excitatory (*ee*) connections over pulses for canonical iTBS, as demonstrated in prior modelling [43].

### Corticothalamic resting-state spectra and component measurement

Resting-state power spectra were computed using Welch’s method from the voltage *V* of the excitatory cortical population using 120 seconds of activity prior to (pre) and following (post) iTBS administration. This population was chosen for its representation of the primary activity captured by EEG. Post-stimulation resting-state activity segments were sampled an additional 100 seconds after iTBS concluded to mitigate residual instability from affecting oscillatory dynamics and to allow for time-delayed *ν* values to converge towards 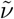 values - with 100 seconds being the discrepancy timescale between signalled and expressed plasticity. Broadband power was computed from the area-under-curve (AUC) of the pre- and post-stimulation resting-state spectra (.25-50 Hz bin range) using the NumPy trapz function, formulated as

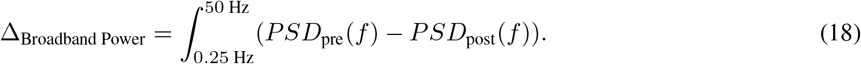

In addition, alpha peak frequency and power were captured using the FOOOF library ([127]). Shifting the circuit’s resting-state alpha frequency between 8-13 Hz was achieved by modifying corticothalamic delay timescales (*t*_0_) from a standard.0425 s (the time for a signal to unidirectionally propagate between cortical and thalamic tracts) to between.055 -.030 s for slower and faster frequencies, respectively. This mechanism is well understood and discussed in [41]. Alpha power change was assessed as the same pre-post difference as broadband power change. Alpha resonance distance (ARD) was computed as the standard deviation of a protocol’s inter-burst frequency from the circuit’s resting-state natural and first subharmonic alpha frequency. Smaller values equate to a smaller discrepancy between stimulation-endogenous frequency alignment.

### Measurement of synaptic weight change

Synaptic weight change (Δ*ν*) were computed as the difference between the absolute post- and pre-stimulation connections weights, where a positive value denotes functional enhancement (further from 0, relative to its initial weight; LTP) and a negative value denotes dampening (closer to 0, relative to its initial weight; LTD). Synaptic weight change was assessed across individual connections and as a circuit-mean coefficient.

### Measurement of calcium volumes

Calcium influx was induced during active iTBS. Calcium volumes were computed as the number of instances wherein stimuli drove calcium concentrations within LTD (0.25 – 0.45 *µ*mol) and LTP (> 0.45 *µ*mol) threshold ranges [45, 44, 43].

### Measurement of loop gain change

Akin to synaptic weight change, corticothalamic loop gain change (see section *Corticothalamic loop gains and space*) was computed as the difference between post and pre-stimulation gains, with positive values denoting functional enhancement and negative values denoting dampening.

### Stimulation intensity and protocol amplitude scaling

Across the parameter space, protocols of greater pulses-per-burst and inter-burst frequencies innately deliver stimuli faster than protocols with smaller parameter values. It was hypothesized that iTBS-induced effects would scale linearly with stimulation intensity, such that protocols that administer greater pulse volumes per train would modulate synaptic weights and oscillatory power more strongly than those that administer fewer. To account for the scaling stimulation intensities across the parameter space, isolate the temporo-modulatory effects of pulse pattern from intensity, and nullify effects of neural adaptation for different stimulation patterns, protocol pulse amplitudes were scaled as a product of their pulse rate (the process of which is depicted in Fig. S5). First, the pulse amplitude of canonical iTBS was established by finding the value that induced a steady-state oscillation in *ee* synaptic weight signaling 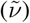, as was first demonstrated in a single-population model by [43]. Once the value was determined, the pulse amplitudes of all other protocols were scaled by their relative intensity to canonical iTBS, such that protocols of lesser relative intensity were administered with greater amplitudes (capped at double the canonical amplitude) and protocols of greater relative intensity were administered with smaller amplitudes, respectively, explicitly calculated as the number of pulses delivered within a single iTBS train:

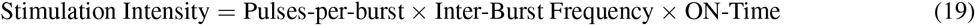

the dot product of the number of pulses-per-burst and the inter-burst frequency, multiplied by the length (in seconds) of the ‘active’ train stimulation period (fixed at a standard 2-second ON). Protocols amplitudes were adjusted through this scaling to produce equivalent stimulation intensities. To further validate the observed effects, supplemental analyses also tested protocols with a fixed amplitude derived from the approach used to determine the canonical iTBS value (see *Supplementary materials*).

### rTMS parameters

rTMS protocols were varied across pulses-per-burst (#) and inter-burst frequency (Hz) parameters. Of the 432 unique protocols, 219 were included for statistical analyses based on their capacity to modulate broadband power at or beyond 10% of the maximum power modulation induced. This was done to ensure a minimal threshold of induced modification was included and non-effect protocols did not confound statistical outcomes.

### Statistical testing

A threshold of p <.001 was used to classify statistical significance. Bonferroni multiple comparison corrections were performed for multiple linear regression analyses of individual alpha frequencies, synaptic weights and calcium LTP and LTD concentrations for each connection, and circuit loop gains.

### Computing

Data loading, processing, analysis, and visualization were computed using Python 3.8.6 on the Compute Canada: Niagara high-performance computing cluster (Digital Research Alliance of Canada) and the CAMH Specialized Computing Cluster (SCC). Neural mass model simulations were performed using the open-source NFTsim C++ package [126]. The codebase for simulating and analyzing this work may be found at: https://github.com/GriffithsLab/Kadak2025_alpha-subharmonics-rtms-eeg.

## Tables

### Alpha Shifts

**Table 1:** MLR coefficients between ARD and broadband power modulation over unique corticothalamic alpha frequencies. Bolded values represent statistical significance following multiple comparison corrections.

| Alpha frequency (Hz) | $t_0/2$ (s) | ARD (IV) |
| --- | --- | --- |
| 13.0 | .0300 | **t = -7.42, p < .001** |
| 12.5 | .0305 | **t = -7.20, p < .001** |
| 12.0 | .0310 | **t = -5.01, p < .001** |
| 11.5 | .0320 | **t = -4.11, p < .001** |
| 11.0 | .0360 | t = -3.88, p < .001 |
| 10.5 | .0425 | **t = -10.01, p < .001** |
| 10.0 | .0460 | **t = -6.18, p < .001** |
| 9.5 | .0470 | **t = -5.98, p < .001** |
| 9.0 | .0480 | **t = -5.45, p < .001** |
| 8.5 | .0490 | **t = -4.47, p < .001** |
| 8.0 | .0550 | t = -1.85, p = .711 |

### Synaptic Weight

**Table 2:** MLR coefficients for iTBS-induced modifications to corticothalamic connection weights (Δ*ν*), predictive of broadband power modulation and predicted by ARD. Bolded values represent statistical significance following multiple comparison corrections.

| Connection | Broadband power (DV) | ARD (IV) |
| --- | --- | --- |
| ee | **t = -6.53, p < 0.001** | **t = 7.22, p < 0.001** |
| ei | **t = -6.23, p < 0.001** | **t = 6.93, p < 0.001** |
| es | **t = -6.17, p < 0.001** | **t = 6.90, p < 0.001** |
| ie | **t = 7.33, p < 0.001** | t = -0.55, p = 0.582 |
| ii | **t = 9.16, p < 0.001** | t = -1.44, p = 0.151 |
| is | **t = 9.15, p < 0.001** | t = -1.43, p = 0.154 |
| re | **t = 25.63, p < 0.001** | **t = -5.81, p < 0.001** |
| rs | **t = 23.64, p < 0.001** | **t = -6.56, p < 0.001** |
| se | t = 2.59, p = 0.010 | t = -2.70, p = 0.007 |
| sr | t = 2.59, p = 0.010 | t = -2.71, p = 0.007 |

### Calcium

**Table 3:** MLR coefficients for induced LTD & LTP calcium volumes, predictive of broadband power modulation and predicted by ARD. Bolded values represent statistical significance following multiple comparison corrections.

| Connection | Broadband power (DV) |  | ARD (IV) |  |
| --- | --- | --- | --- | --- |
|  | LTD | LTP | LTD | LTP |
| ee | **t = 12.56, p < 0.001** | t = 2.85, p = 0.005 | t = -3.38, p = 0.001 | t = 3.32, p = 0.001 |
| ei | **t = 10.73, p < 0.001** | t = 0.47, p = 0.636 | -3.22, p = 0.001 | **t = 4.68, p < 0.001** |
| es | **t = 10.64, p < 0.001** | t = 0.44, p = 0.660 | -3.20, p = 0.002 | **t = 4.70, p < 0.001** |
| ie | t = 0.58, p = 0.564 | **t = 8.64, p < 0.001** | t = 1.54, p = 0.126 | t = 0.01, p = 0.996 |
| ii | t = 0.39, p = 0.693 | **t = 9.69, p < 0.001** | t = 2.53, p = 0.012 | t = -0.40, p = 0.689 |
| is | t = 0.40, p = 0.689 | **t = 9.69, p < 0.001** | t = 2.50, p = 0.013 | t = -0.40, p = 0.687 |
| re | t = 1.39, p = 0.167 | **t = 17.42, p < 0.001** | t = 4.09, p < 0.001 | t = -3.87, p < 0.001 |
| rs | t = 1.57, p = 0.118 | **t = 16.20, p < 0.001** | 4.02, p < 0.001 | **t = -5.75, p < 0.001** |
| se | t = -2.21, p = 0.028 | N/A | 2.15, p = 0.033 | N/A |
| sr | t = -2.21, p = 0.028 | N/A | 2.15, p = 0.033 | N/A |

### Circuit loop gains

**Table 4:** MLR coefficients of corticothalamic loop gains change, predictive of broadband power modulation and predicted by ARD. Bolded values represent statistical significance following multiple comparison corrections.

| Loop | Broadband power (DV) | ARD (IV) |
| --- | --- | --- |
| X | **t = 4.18, p < 0.001** | t = 2.02, p = 0.045 |
| Y | **t = -23.05, p < 0.001** | **t = 5.74, p < 0.001** |
| Z | **t = 25.46, p < 0.001** | **t = -8.22, p < 0.001** |

### Model parameters

**Table 5:**
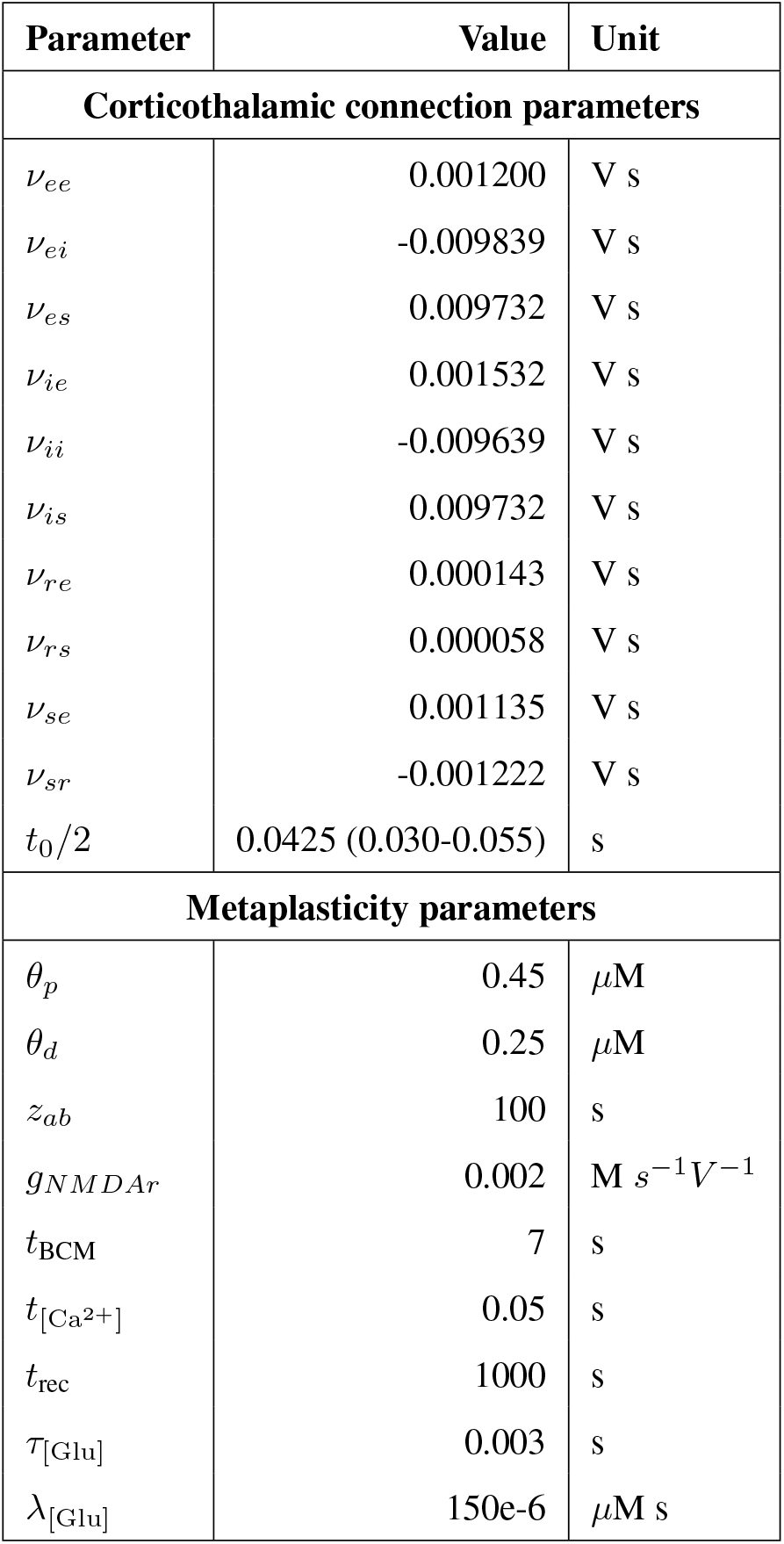
Corticothalamic neural population with metaplasticity model parameters.

## Supplementary materials

Here we provide a set of additional content for rTMS-induced changes of individual synaptic weights and circuit loop gains, as well as alpha-band power changes and their relationship with said outcomes.

**Figure S1:**
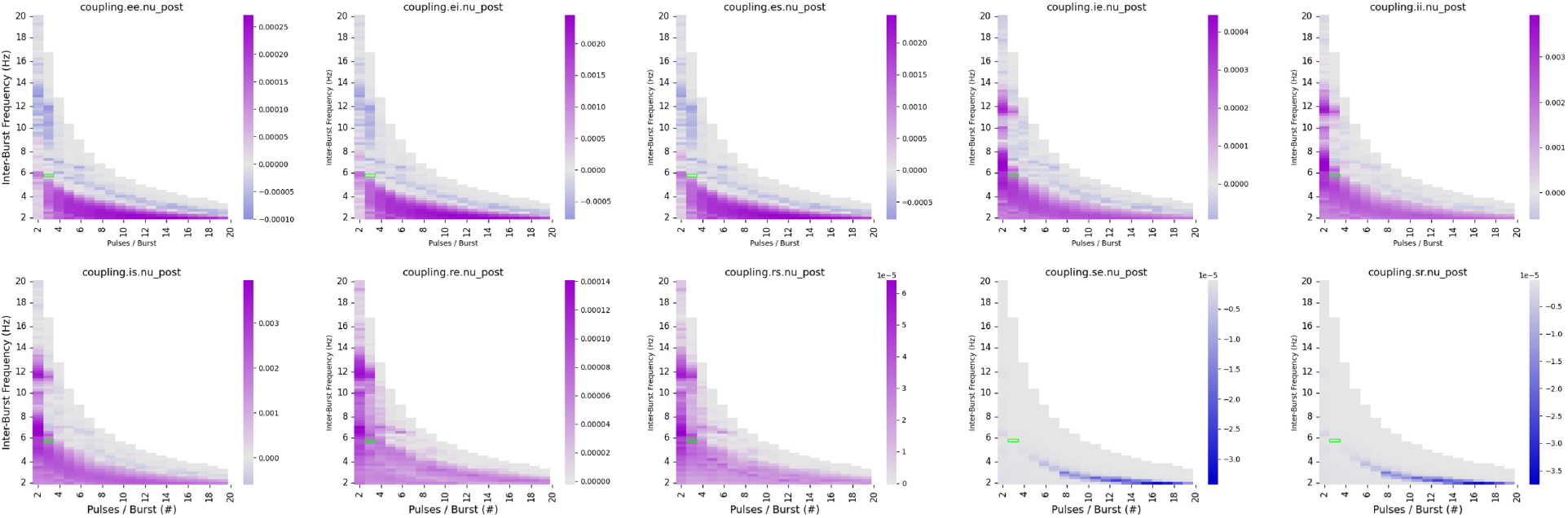
Heatmap of post-iTBS connection weights for each connection within the corticothalamic circuit. Purple denotes weight enhancements, while blue denotes reductions.

**Figure S2:**
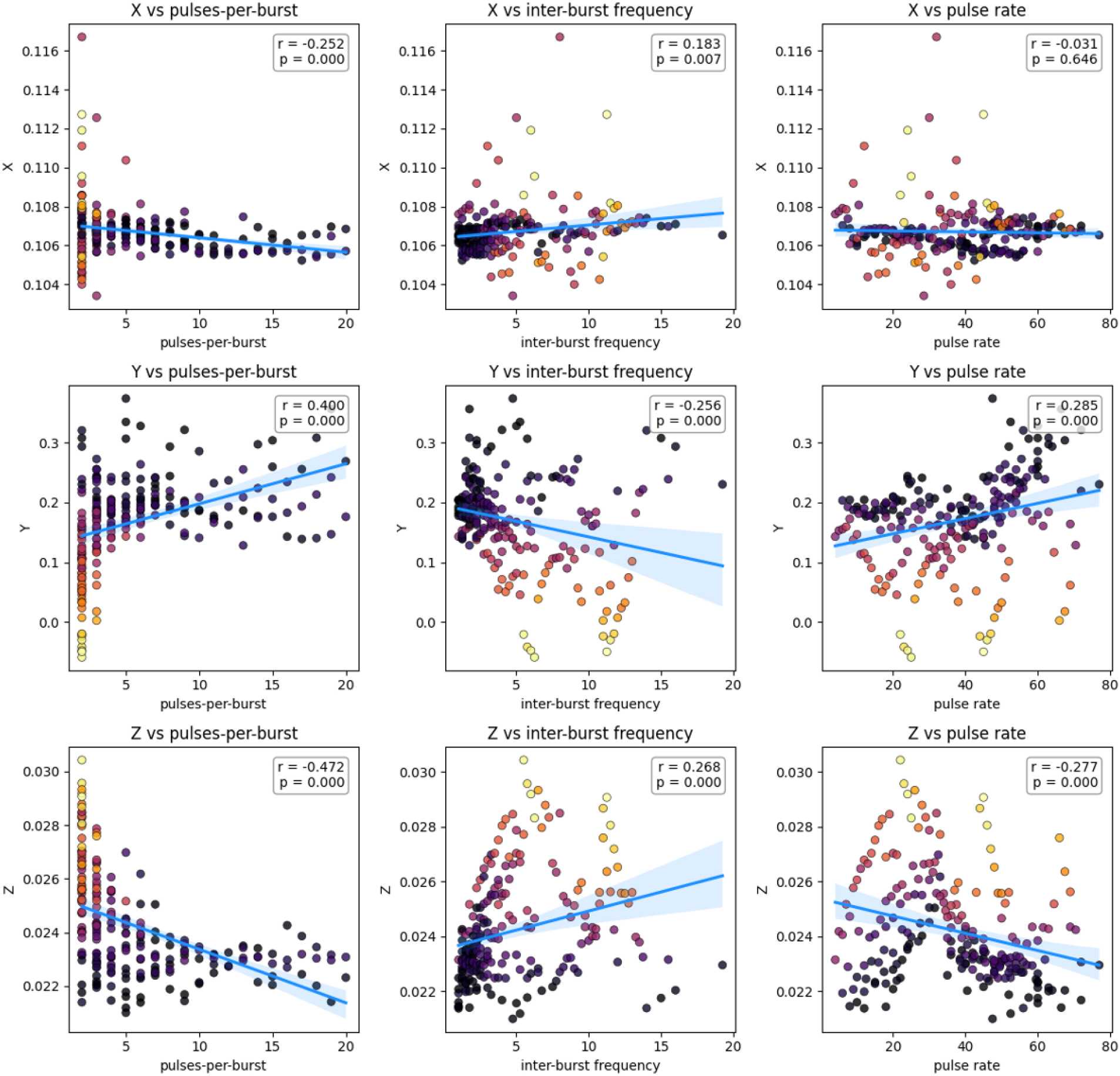
Scatterplot distributions of post-stimulation X, Y, and Z loop gains across pulses-per-burst, inter-burst frequency, and pulse rate iTBS parameters.

**Figure S3:**
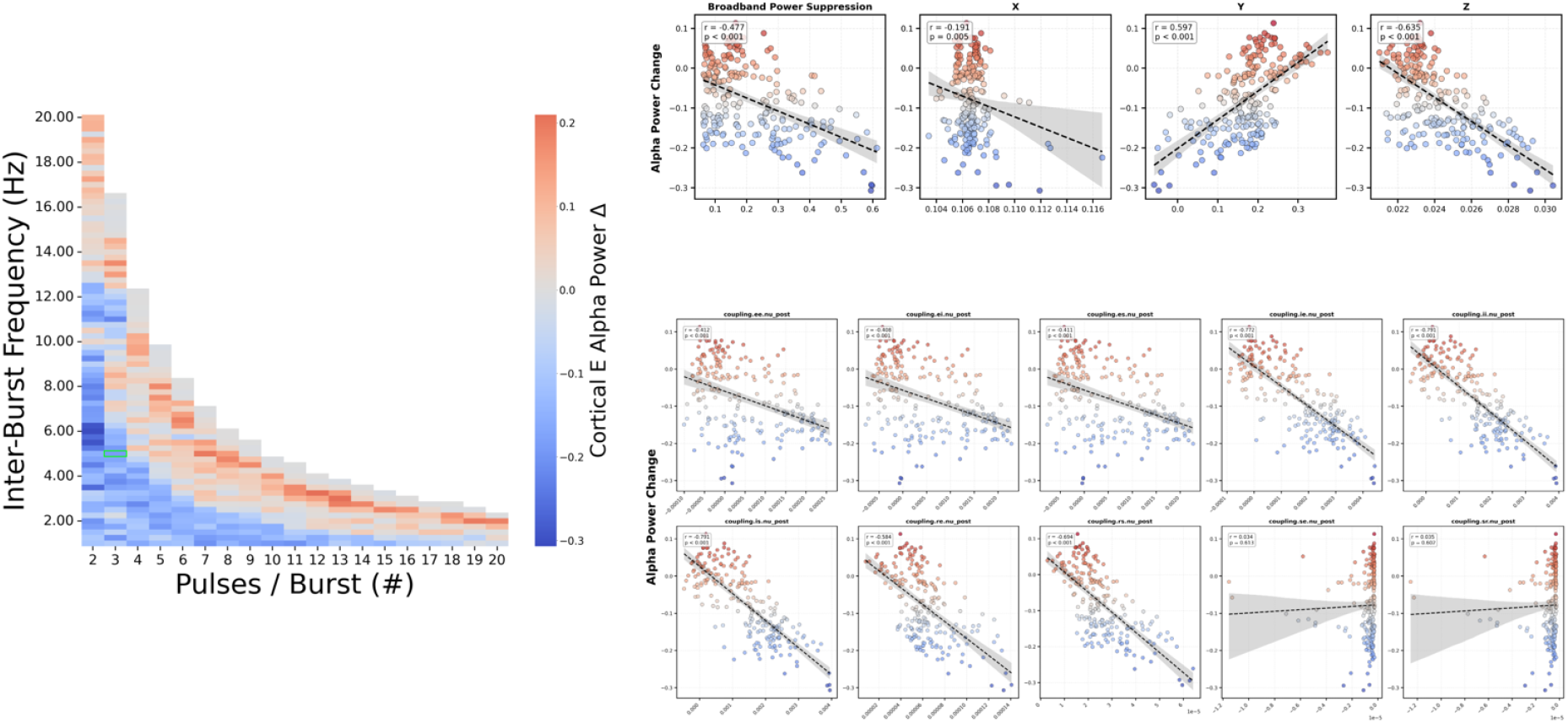
Relationship between iTBS-induced changes of alpha and broadband power, connection weights, and loop gains.

### Scaled vs. fixed metaplastic effects

We hypothesized that protocols that avoid oversaturating NMDAR conductance rates would drive a more consistent calcium influx and induce their effects more efficiently than those that cause fluctuations in conductance rates and subsequent calcium levels. We aimed to isolate the stimulation effects of pulse pattern and frequency from the byproduct of intensity by scaling pulse amplitudes in proportion to their administered rate, such that faster protocols had smaller amplitudes (see *Stimulation intensity and protocol amplitude scaling*). The results showed that the influence of protocol intensity on outcome measures was indeed removed, allowing us to focus on the patterned effects of stimuli. Only fixed-amplitude protocols demonstrated a robust significant relationship between pulse rate and engagement of circuit-mean metaplastic homeostasis (quantified as the standard deviation of NMDAR conductance rates during active iTBS). Additionally, metaplastic homeostasis was negatively related to broadband power modulation and plasticity-inducing calcium levels, and positively related to circuit-wide synaptic weight change. These effects were largely absent in scaled-amplitude protocols, with only circuit-mean connection weight change yielding a significant relationship NMDAR conductance rates (Supplemental Fig. S7). While the extent of connection weight change differed, the modification patterns associated with treatment efficacy were persistent between conditions. As depicted in the supplemental radar plots (Supplemental Fig. S6), there was a strong similarity in the relative synaptic modification patterns between scaled (purple) and fixed (green) simulations, validating that the induced effects are consistent between conditions.

These trends demarcate our scaled and fixed-amplitude conditions and confirm that our correction approach effectively ‘detrended’ the scaling energy contribution over pulse rate to isolate the influence of pulse pattern outcome measures. It is important to note these NMDAR metaplasticity mechanisms were still very much engaged and functionally relevant in system dynamics and measured outcomes, but that the extent of their engagement was systematically corrected to enable more physiologically plausible and experimentally practical examinations of the parameter space. Our approach emphasizes that balancing pulse pattern and intensity, and understanding the role of metaplastic homeostasis mediating their effects, is crucial for formulating effective rTMS protocols.

We discern that protocols minimizing the engagement of metaplastic NMDAR homeostasis facilitate a more consistent calcium influx, leading to stronger and more efficient modulatory effects. Conversely, protocols that engage metaplastic homeostasis more aggressively tend to result in fluctuating calcium permeability and less efficient plasticity trends.

**Figure S4:**
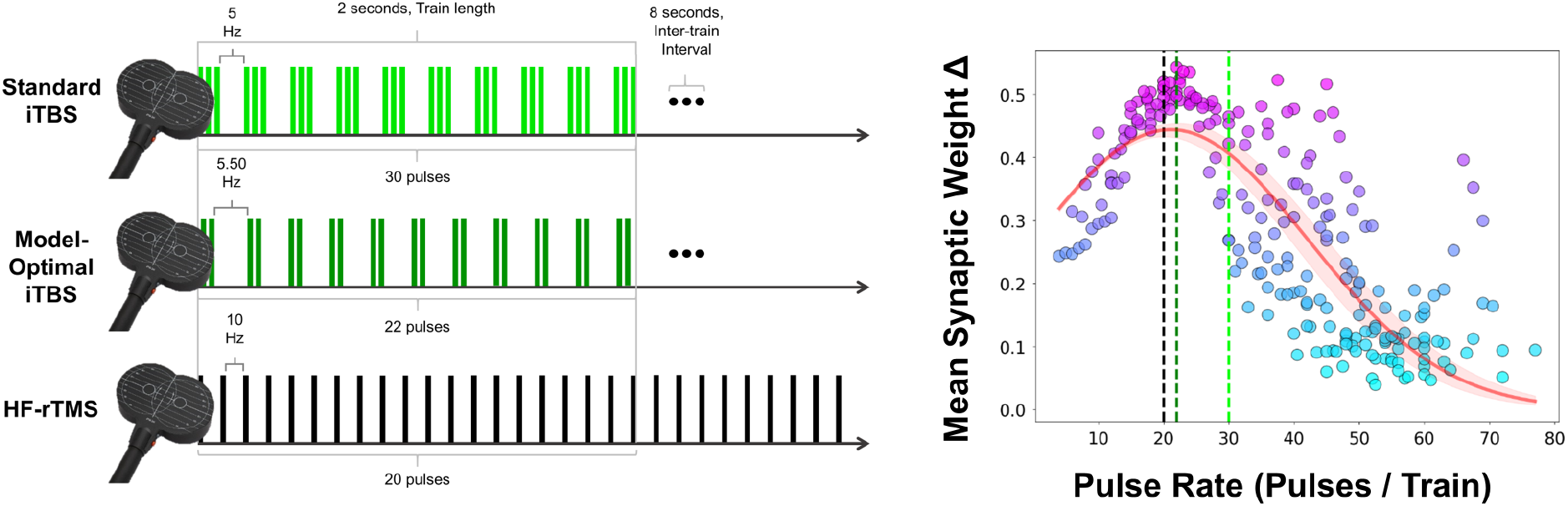
Illustration of circuit-mean synaptic weight change across pulse rate. Circuit-mean synaptic weight change over pulse rates, featuring canonical iTBS (lime), model-optimal iTBS (dark green), and HF-rTMS (black).

**Figure S5:**
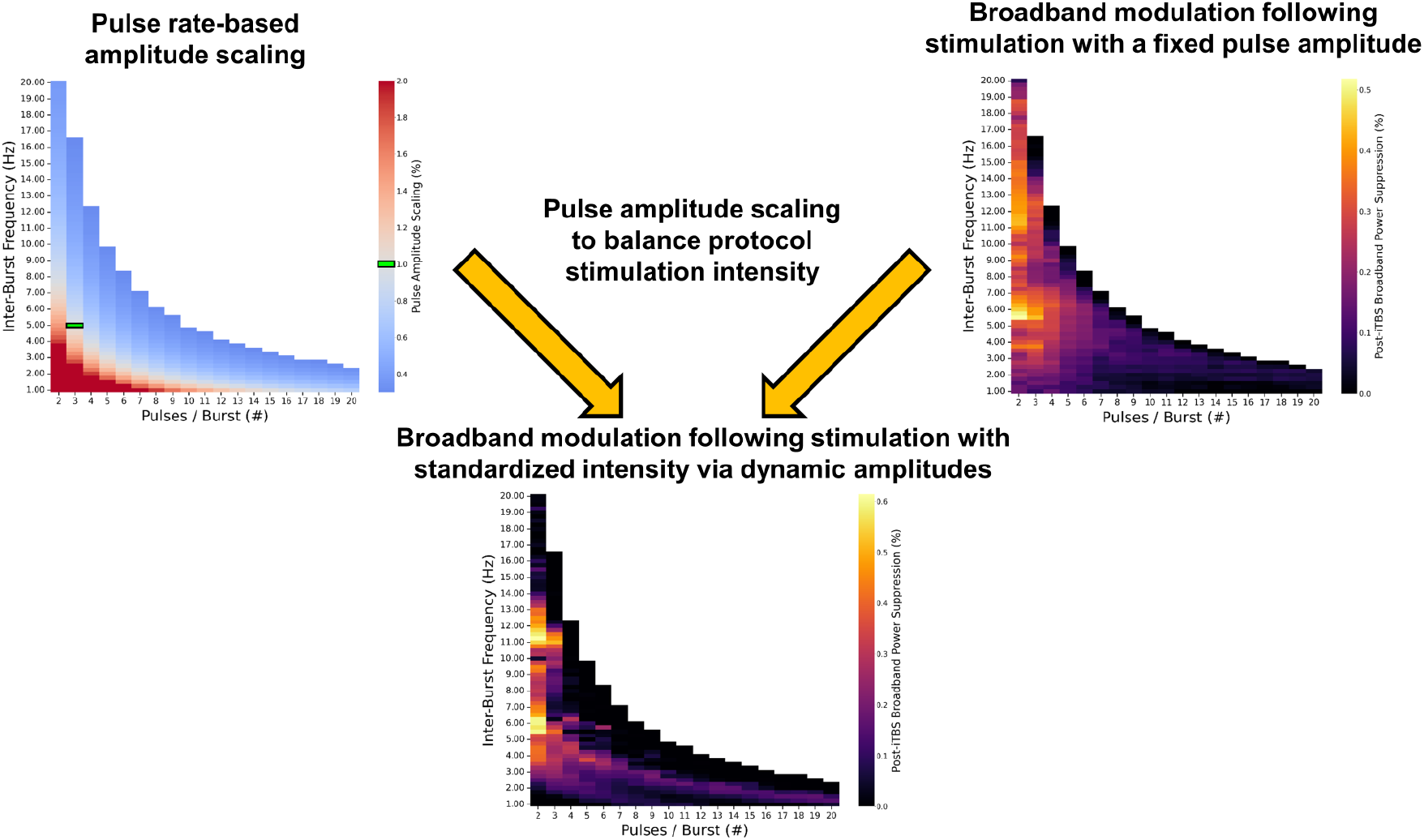
Method for standardizing protocol intensity. Scaling is based on the proportional stimulation intensity of canonical iTBS.

**Figure S6:**
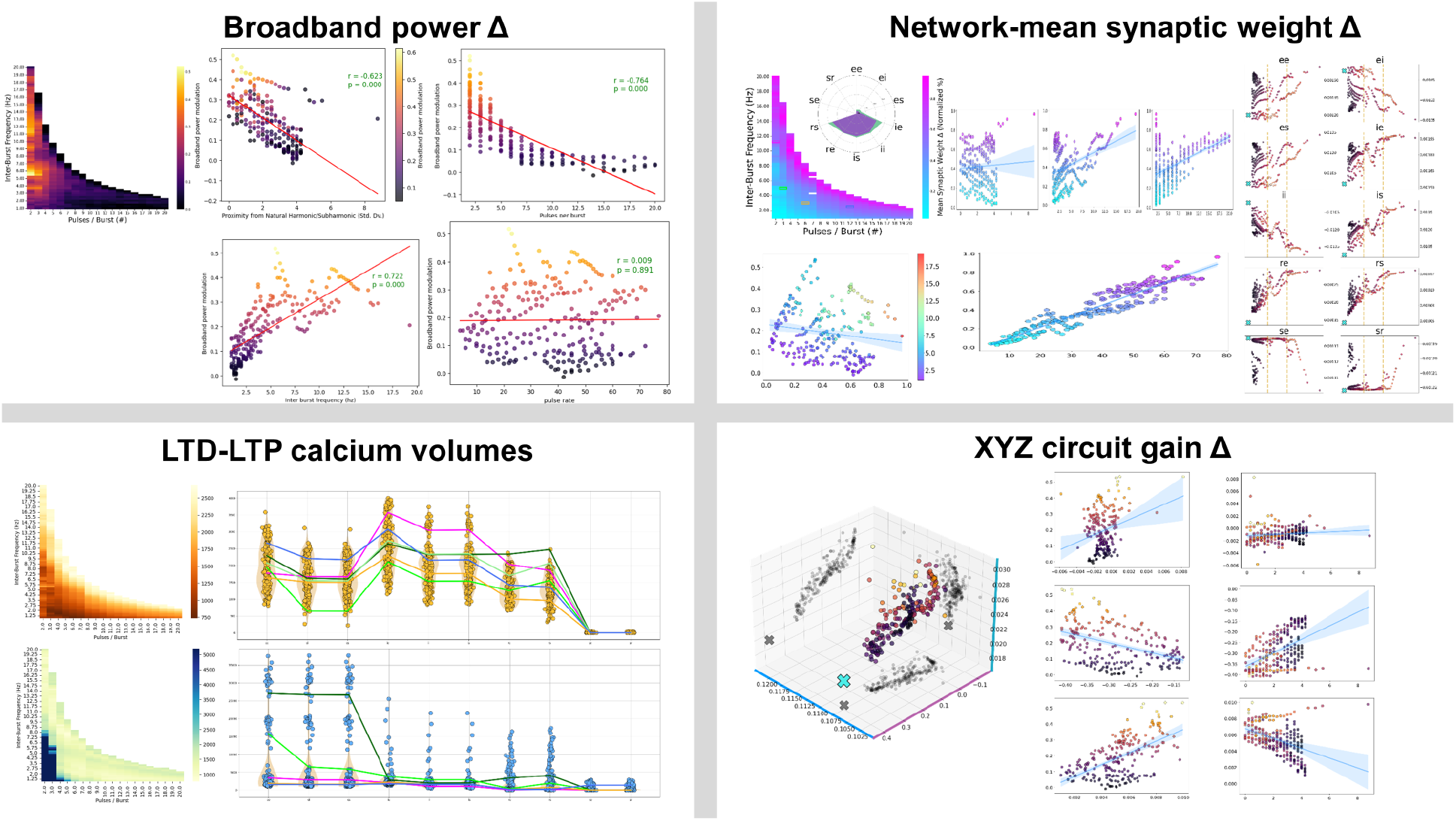
Primary analyses conducted with protocols using a fixed pulse amplitude. rTMS-induced effects on broadband power modulation, Circuit-mean and individual synaptic weight change, LTD and LTP calcium levels, and X, Y, and Z loop gains.

**Figure S7:**
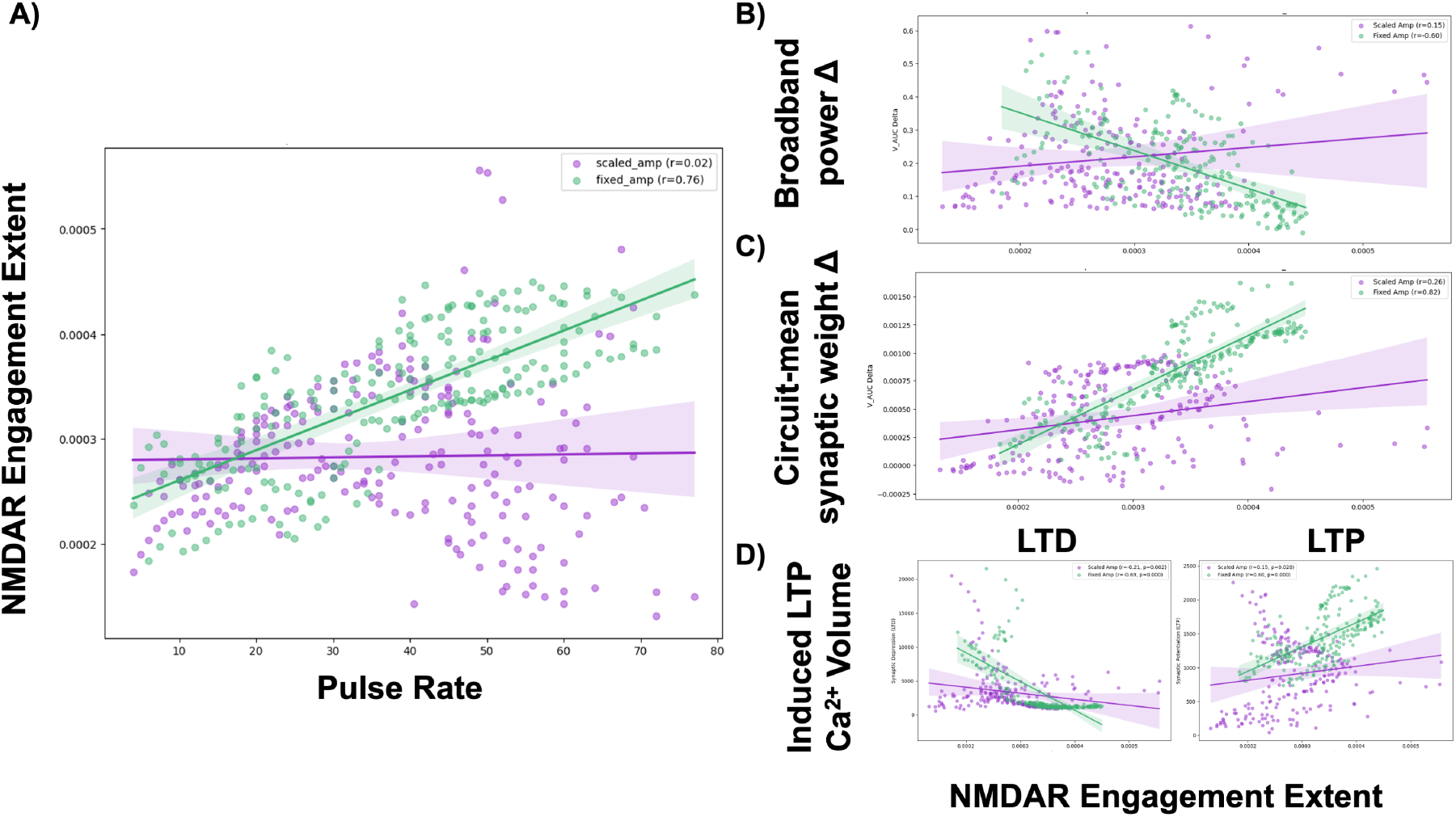
Comparison of active-stimulation NMDAR conductance variance between scaled- (purple) and fixed- (green) amplitude iTBS protocols. **A)** Relationship between protocol pulse rate and standard deviation of NMDAR conductance. Relationship between NMDAR engagement and **B)** broadband power modulation, **C)** circuit-mean synaptic weight change, and **D)** plasticity-inducing calcium volumes.

